# Transcranial direct current stimulation alters cerebrospinal fluid-interstitial fluid exchange in mouse brain

**DOI:** 10.1101/2023.12.30.573695

**Authors:** Yan Wang, Hiromu Monai

**Author notes:** To whom correspondence should be addressed: Hiromu Monai. **Conflict of interest statement:** No COI.

## Abstract

**Background:** Transcranial direct current stimulation (tDCS) is a non-invasive brain stimulation technique that has gained prominence recently. Clinical studies have explored tDCS as an adjunct to neurologic disease rehabilitation, with evidence suggesting its potential in modulating brain clearance mechanisms. The glymphatic system, a proposed brain waste clearance system, posits that cerebrospinal fluid-interstitial fluid (CSF-ISF) exchange aids in efficient metabolic waste removal. While some studies have linked tDCS to astrocytes inositol trisphosphate (IP_3_)/Ca^2+^ signaling, the impact of tDCS on CSF-ISF exchange dynamics remains unclear.

**Hypothesis:** tDCS influences the dynamics of CSF-ISF exchange through astrocytic IP_3_/Ca^2+^ signaling.

**Methods:** In this study, we administered tDCS (0.1mA for 10 minutes) to C57BL/6 mice anesthetized with ketamine-xylazine (KX). The anode was positioned on the cranial bone above the cortex, and the cathode was inserted into the neck. Following tDCS, we directly assessed brain fluid dynamics by injecting biotinylated dextran amine (BDA) as a CSF tracer into the cisterna magna (CM). The brain was then extracted after either 30 or 60 minutes and fixed. After 24 hours, the sectioned brain slices were stained with Alexa 594-conjugated streptavidin (SA) to visualize BDA using immunohistochemistry. We conducted Electroencephalography (EEG) recordings and aquaporin 4 (AQP4)/CD31 immunostaining to investigate the underlying mechanisms of tDCS. Additionally, we monitored the efflux of Evans blue, injected into the cisterna magna, using cervical lymph node imaging. The experiments were subsequently repeated with inositol trisphosphate receptor type 2 (IP_3_R2)-knockout mice.

**Results:** Post-tDCS, we observed an increased CSF tracer influx, indicating a modulation of CSF-ISF exchange by tDCS. Additionally, tDCS appeared to enhance the brain’s metabolic waste efflux. EEG recordings showed an increase in delta wave post-tDCS. But no significant change in AQP4 expression was detected 30 minutes post-tDCS.

**Conclusion:** Our findings suggest that tDCS augments the glymphatic system’s influx and efflux. Through astrocyte IP_3_/Ca^2+^ signaling, tDCS was found to modify the delta wave, which correlates positively with brain clearance. This study underscores the potential of tDCS in modulating brain metabolic waste clearance.

## Introduction

Non-invasive brain stimulation (NIBS), which includes transcranial magnetic stimulation (TMS) and transcranial electric stimulation (such as transcranial direct current stimulation or tDCS), has gained increasing interest in treating neurological diseases [1–3]. NIBS offers localized regulation of brain neural excitability with typically mild side effects [4–8], presenting a significant advantage over drug therapies that often have severe adverse events. Consequently, NIBS holds considerable potential for diagnosis, treatment, and rehabilitation of neurological conditions.

Transcranial direct current stimulation (tDCS), a form of NIBS, is under investigation in numerous clinical trials targeting neurological diseases. Yet, it has not received U.S. Food and Drug Administration (FDA) approval for clinical use [9,10]. tDCS involves positioning target and reference electrodes on the scalp to administer a low direct current (1-2 mA) for 10-30 minutes [11,12]. It has been explored as a complementary therapy in clinical studies for various neurological disorders [2,13–16]. For instance, a clinical trial addressing Alzheimer’s disease reported that tDCS increased the serum levels of Aβ 1-42, potentially diminishing its brain accumulation [14]. Preclinical research has shown that tDCS can reduce Aβ in Alzheimer’s model mice [17], hinting at its influence on brain clearance mechanisms.

The concept of the glymphatic system, a brain metabolic waste clearance pathway, was recently introduced by Iliff et al. [18]. This system begins with the production of cerebrospinal fluid (CSF) in the choroid plexuses and possibly other extrachoroidal sites. The CSF then enters and circulates through periarterial spaces into the brain parenchyma, exchanges with the interstitial fluid (ISF), and finally exits the brain via lymphatic vessels and alternative routes [19,20]. Factors such as sleep and anesthesia are known to facilitate the removal of brain waste, including Aβ [21]. The water channel protein aquaporin 4 (AQP4), located at the endfeet of perivascular astrocytes, is crucial for CSF-ISF exchange [18,22]. However, the impact of tDCS on the glymphatic system’s function remains to be elucidated. In this study, we employed immunohistochemistry to directly visualize changes in brain fluid dynamism and CSF-ISF exchange post-tDCS.

## Materials and Methods

All experimental protocols were approved by the Institutional Animal Care and Use Committee of Ochanomizu University, Japan (animal study protocols 22017). All animal experiments were performed according to the guidelines for animal experimentation of Ochanomizu University that conforms with the Fundamental Guidelines for Proper Conduct of Animal Experiment and Related Activities in Academic Research Institutions (Ministry of Education, Culture, Sports, Science and Technology, Japan). Efforts were taken to minimize the number of animals used. This study was carried out in compliance with the ARRIVE guidelines.

### Animals

Adult male and female C57BL/6, and type2 IP_3_ receptor knock-out (IP_3_R2 KO) mice were used (aged 8-12 weeks, weight between 20 g and 25 g). Mice were housed under a 12 h/12 h light/dark cycle and raised in groups of up to five mice. IP_3_R2 KO mice are available from the RIKEN BioResource Center (Resource ID: RBRC10289).

### Anesthesia

Mice were anesthetized by cocktail of ketamine and xylazine (KX, 70 mg kg^−1^ ketamine and 10 mg kg^−1^ xylazine) via intraperitoneal injection. Half the anesthetic dose was replenished every 30 min for C57BL/6 mice and 60 min for IP_3_R2 KO mice, and the mice were always observed for their anesthetic state by their heart rate and respiration rate.

### Transcranial direct current stimulation application

The mice were anesthetized and fixed on a head holder with the head held horizontally, the scalp was cut off, and a conductive gel (1.5% agarose in phosphate buffered saline, PBS) with a 1 mm radius was placed over the skull (2.00 mm posterior, 2.00 mm lateral from bregma), then the positive electrode was applied, the skin over the neck was cut off, and the negative electrode was inserted into the muscle of the neck for stimulation (0.1 mA, 10 min).

### Cisterna magna injection

After anesthesia, mice were immobilized on a head holder with the head down and at an angle of 120 degrees to the body. The skin above the cisterna magna (CM) was cut open, and the muscle was grasped with forceps to expose the CM. A glass tube with the cerebrospinal fluid (CSF) tracer, biotinylated dextran amines (BDA, 70 kMW, D1957) was inserted into the CM using a micromanipulator (NARISHIGE, MDS-1), and 1% CSF tracer was injected into the CM with a syringe pump (1 μl/min, 5 μl). After the injection, the mice were kept in a prone position on a heating pad. After 30 or 60 min, the brains were removed and stored in 4% paraformaldehyde solution (PFA) for 24 h in a refrigerator.

### Streptavidin staining

The brains obtained after CM injection were cut into 60 μm slices using a micro slicer (DOSAKA, DTK-1000N). Then the BDA was visualized by staining with streptavidin (SA, S32356, 1:1000 in 0.1% phosphate-buffered saline with Triton (PBST)) at room temperature for one hour, after which they were washed three times with PBS and made into microscope samples. Images were taken with the microscope (OLYMPUS, BX53F2, 4X objective lens, N5702800) and analyzed with ImageJ.

### In vivo cervical lymph node imaging

After the mice were anesthetized, the neck was depilated with depilatory cream, then the skin was cut, and cervical lymph nodes were exposed. Mice were placed under the microscope for the pre-injection images. Direct Blue 53 (DB53 or Evans blue, 1%, 960 MW, E2129, 5 μl in total) was injected via CM injection, and mice were placed on a heating pad. After injection, mice were fixed to keep them supine and placed under the fluorescence stereo microscope (EVIDENT, MVX10). The imaging started 30 minutes from the end of the injection.

### Immunohistochemistry

Mice were perfused with PBS and then fixed with 4% PFA perfusion via heart, and then the brains were removed and placed in 4% PFA overnight at 4°C. The brains of the mice were cut into 60 μm slices using a micro slicer and placed in PBS. Then, the brain slices were placed in a blocking solution of 10% Normal goat serum at room temperature, 100 rmp, for 1h, and then replaced with primary antibody (AQP4, A5917, 1:2000; CD31, 550274, 1:1000) at 4°C, 100 rmp, overnight. The samples were washed three times with 0.1% PBST solution, replaced with secondary antibody (Alexa fluor 488, ab150077, 1:1000; Alexa fluor 594, A11007, 1:1000), room temperature, 100 rmp, 1h, washed three times with 0.1% PBST solution and twice with phosphate buffer (PB) solution to make samples for microscopic observation. Images were taken with a confocal microscope (LSM700, ZEISS, Plan-Apochromat 40x/1.4 Oil DIC M27 (FWD = 0.13 mm)) and then analyzed with ImageJ and MATLAB.

### Electroencephalography recordings

After anesthesia, the mice were fixed in a head holder with the head held horizontally, the scalp was cut, holes were punched over the motor area (2.00 mm anterior and 2.0 mm lateral from bregma), and above the cerebellum, positive and negative electrodes were inserted separately. A signal amplifier (HAS-4 Head Amplifier System, Bio Research Center Co. Ltd., Tokyo., Japan) was connected. Data were amplified using head stage (×1000, filtered 0.1–1000, Hz) and recording data was digitized LabVIEW (National Instruments).

### Image processing

For all the images, the channels were split by ImageJ. For sagittal slices, we obtained the mean intensity of each region of interest (ROI, Fig. 1B) for the red channel. For coronal slices, the median intensity of a 250 nm thick cortex starting from the brain’s edge and extending from the highest point on the left to the far left was obtained for the red channel and then averaged across all brain slices for each brain. For in vivo cervical lymph node imaging, the mean intensity of cervical lymph nodes was taken for the red and green channels respectively, and the mean intensity of the red channel was normalized by the green channel to eliminate the interference of autofluorescence. For AQP4/CD31 images, using MATLAB, the location of pixels whose brightness exceeded a threshold in images stained with CD31 was defined as perivascular or peri membranous for ± 5 pixels around them, and neuropil for the rest of the pixels. Analyses were performed using the imbinarize function of MATLAB. The AQP4 in each region was defined as perivascular. The number of pixels in each region in which the AQP4 brightness exceeded the threshold was counted. This result was normalized by the number of pixels exceeding the CD31 threshold.

**Fig.1.**
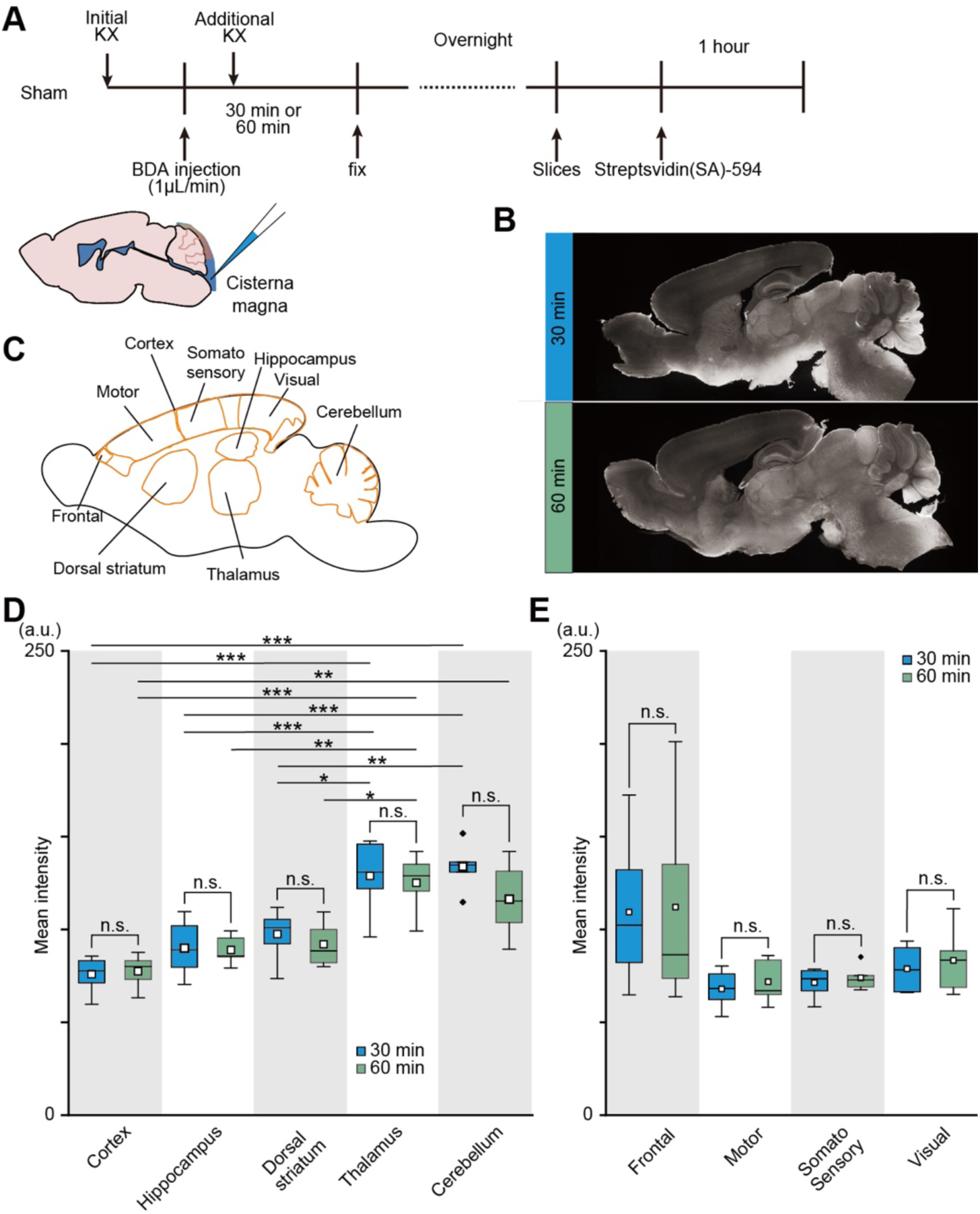
No changes in the CSF tracer after 30 min and 60 min at the end of CM injection in sham mice. A. Schematic diagram for CM injection of CSF tracer. B. Representative images of CSF tracer distribution in sham mice. C. ROIs of data analysis. D-E. The mean intensity of the CSF tracer after 30 min (blue, N = 6) and 60 min (green, N = 5) at the end of CM injection. *p < 0.05, **p < 0.01, ***p < 0.01, two-way ANOVA, Tukey’s post analysis

### Statistical analysis

Two-way analysis of variance (ANOVA) followed by Tukey’s multiple comparisons test was used to compare the 30 min group and the 60 min group for the influx of sham mice. Two-way repeated measures ANOVA followed by Tukey’s multiple comparisons test was used to compare two experimental groups for the different periods of EEG recordings. Student’s t-test was used for other experiments to compare the sham and tDCS groups. All data are expressed as mean±s.e.m.

## Results

### Observed CSF-ISF exchange by a tracer injected from cisterna magna

We initially observed the exchange between cerebrospinal fluid (CSF) and interstitial fluid (ISF) using a tracer to study the clearance of metabolic waste in the brain. Various methods exist for observing cerebrospinal fluid flow, including phase-contrast MRI and radiotracer techniques [24–26]. In our study, we injected biotinylated dextran amine (BDA, 80 kDa) via the cisterna magna (CM) as the CSF tracer into ketamin-xylazine (KX) anesthetized mice. Subsequently, fixed brain slices were stained with fluorescence-tagged streptavidin (SA) (see Methods, **Fig. 1A**). The brains of mice in the unstimulated group (Sham) were examined at both 30 minutes and 60 minutes post-injection (**Fig. 1B**). We found no significant difference in tracer distribution between the 30-minute and 60-minute time points in regions such as the cortex, hippocampus, dorsal striatum, thalamus, and cerebellum (**Figs. 1C-D**, Cortex: 30 min vs. 60 min, 75.91 ± 3.82 vs. 77.49 ± 4.26, p = 0.79; Hippocampus: 30 min vs. 60 min, 89.94 ± 5.83 vs. 88.99 ± 3.64, p = 0.89; Dorsal Striatum: 30 min vs. 60 min, 97.57 ± 5.46 vs. 91.98 ± 5.58, p = 0.49; Thalamus: 30 min vs. 60 min, 128.89 ± 7.69 vs. 125.18 ± 7.39, p = 0.74; Cerebellum: 30 min vs. 60 min, 133.85 ± 4.83 vs. 116.34 ± 9.40, p = 0.15). These results suggested that in the brain under KX anesthesia, adequate CSF-ISF exchange occurs within 30 minutes, and by 60 minutes, saturation is already achieved. Nevertheless, differences in baseline were observed in various brain regions. Notably, higher intensities were observed in the thalamus and cerebellum compared to the cortex, hippocampus, and striatum (**Figs. 1C-D**, 30 min, Cortex vs. Hippocampus vs. Dorsal striatum vs Thalamus vs. Cerebellum: 75.91 ± 3.82 vs. 89.94 ± 5.83 vs. 97.57 ± 5.46 vs. 128.89 ± 7.69 vs. 133.85 ± 4.83; 60 min, Cortex vs. Hippocampus vs.

Dorsal striatum vs Thalamus vs. cerebellum: 77.49 ± 4.26 vs. 88.99 ± 3.64 vs 91.98 ± 5.58 vs. 125.18 ± 7.39 vs. 116.34 ± 9.40). This observation held for the frontal, motor, sensory, and visual regions in sagittal slices (**Fig. 1C**). Among them, particularly high intensity was observed in the frontal cortex (**Fig. 1E**, Frontal: 30 min vs. 60 min, 109.35 ± 15.75 vs. 112.05 ± 25.40, p = 0.93; Motor: 30 min vs. 60 min, 68.05 ± 4.08 vs. 71.91 ± 5.45, p = 0.59; Somatosensory: 30 min vs. 60 min, 71.45 ± 3.14 vs. 73.97 ± 3.11, p = 0.58; Visual: 30 min vs. 60 min, 78.81 ± 4.75 vs. 83.40 ± 8.20, p = 0.64).

### tDCS increased the CSF-ISF influx

To investigate the potential impact of transcranial direct current stimulation (tDCS) on the exchange between cerebral spinal fluid (CSF) and interstitial fluid (ISF), we administered biotinylated dextran amine (BDA) as a representative CSF tracer. This was done intrathecally at a dose of 10 μl per mouse 30 minutes after a 10-minute tDCS session (**Figs. 2A-B**). Following BDA administration, mouse brains were harvested and analyzed at two distinct time points: 30 minutes and 60 minutes post-injection (**Figs. 2A-B**). Sagittal brain slices were prepared and stained with fluorescence-tagged streptavidin (SA) to visualize BDA distribution (**Fig. 2B**). Our statistical analysis indicated a significant increase in CSF tracers in the cortex, hippocampus, and dorsal striatum 30 minutes after injection post-tDCS (**Fig. 2C**, Cortex: Sham vs. tDCS, 75.91 ± 3.82 vs. 95.37 ± 2.66, p = 2.25E-3; Hippocampus: Sham vs. tDCS, 89.94 ± 5.83 vs. 112.45 ± 4.65, p = 1.29E-2; Dorsal striatum: Sham vs. tDCS, 97.57 ± 5.46 vs. 114.00 ± 2.59, p = 2.90E-2). This increase was particularly pronounced in the motor, somatosensory, and visual regions of the cerebral cortex (**Fig. 2D**, Motor: Sham vs. tDCS, 68.05 ± 4.08 vs. 89.15 ± 3.08, p = 2.18E-3; Somatosensory: Sham vs. tDCS, 71.45 ± 3.14 vs. 92.18 ± 3.22, p = 7.60E-4; Visual: Sham vs. tDCS, 78.81 ± 4.75 vs. 95.59 ± 4.37, p = 2.53E-2). By the 60-minute mark, although CSF tracer levels continued to rise in the aforementioned regions, the somatosensory region of the cortex exhibited a notably higher increase (**Figs. 2E-F**, Cortex: Sham vs. tDCS, 77.49 ± 4.26 vs. 91.56 ± 3.91, p = 3.70E-2; Hippocampus: Sham vs. tDCS, 88.99 ± 3.64 vs. 103.37 ± 4.94, p = 4.12E-2; Dorsal striatum: Sham vs. tDCS, 91.98 ± 5.58 vs. 117.86 ± 5.85, p = 9.77E-3; Somatosensory: Sham vs. tDCS, 73.97 ± 3.11 vs. 88.98 ± 3.93, p = 1.35E-2).

**Fig. 2.**
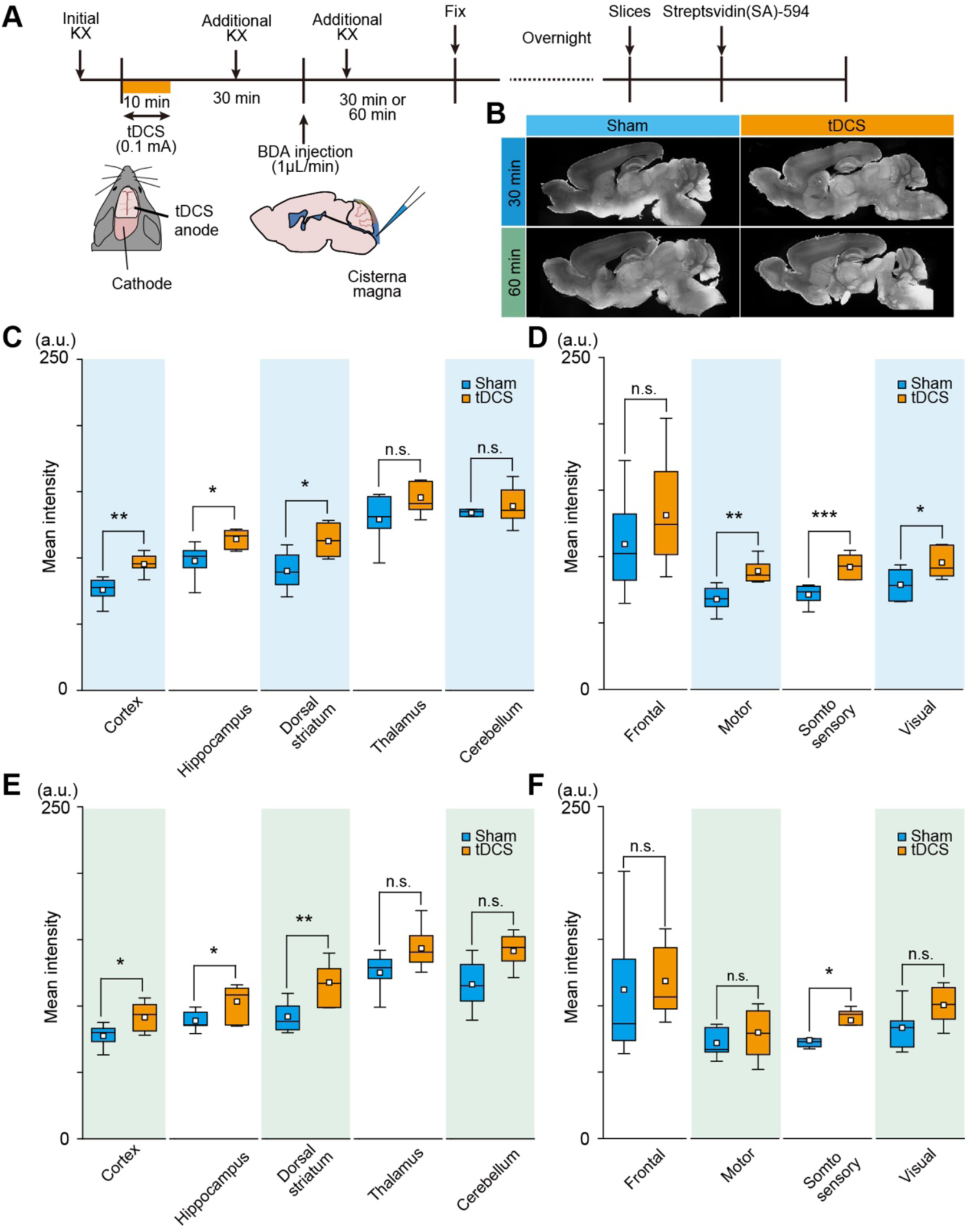
CSF tracer increased in sagittal slices after tDCS. A. Schematic diagram for tDCS and the experiment schedule. B. Representative images of CSF tracer distribution in sagittal slices. C-D. The mean intensity of the CSF tracer after 30 min at the end of CM injection in sagittal slices for sham group (blue, N = 6) and tDCS group (orange, N = 7). E-F. The mean intensity of the CSF tracer after 60 min at the end of CM injection in sagittal slices for sham group (blue, N = 5) and tDCS group (orange, N = 7). *p < 0.05, **p < 0.01, ***p < 0.01, t-test

Coronal brain slices were also prepared and stained with SA (**Fig. 3**). The CSF tracer levels were significantly higher in the tDCS group mice 30 minutes post-injection compared to the sham group (**Fig. 3A-B**, 30 min: Sham vs. tDCS, 106.23 ± 4.40 vs. 121.15 ± 4.66, p = 4.83E-2, **Fig. 3C-D**, 60 min: Sham vs. tDCS, 130.10 ± 12.30 vs. 141.19 ± 15.39, p = 0.59). These observations suggested that tDCS modulates the CSF-ISF exchange dynamics in the mouse brain.

**Fig. 3.**
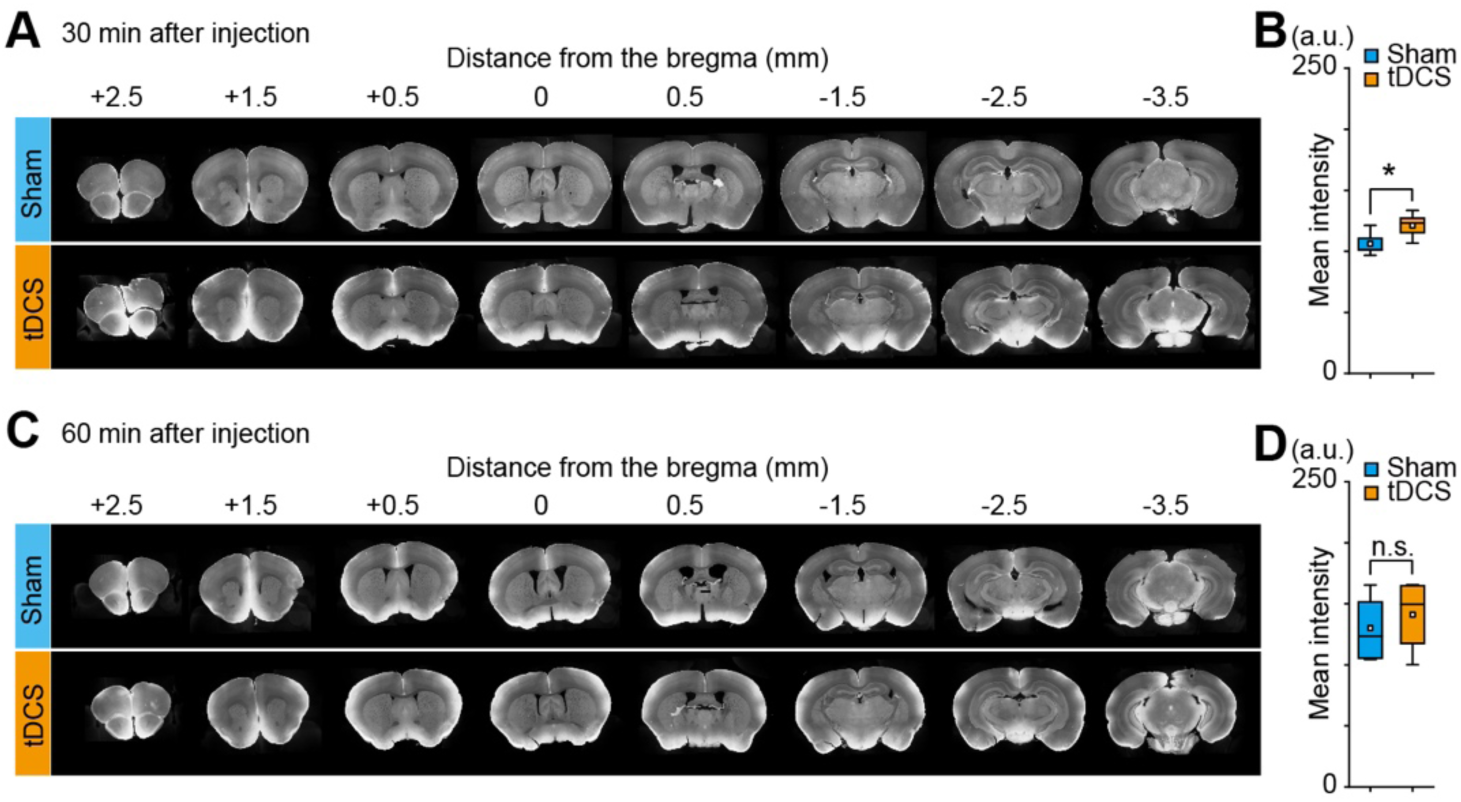
CSF tracer increased in coronal slices after tDCS. A. Representative images of CSF tracer distribution in coronal slices after 30 min at the end of CM injection. B. The mean intensity of CSF tracer in 8 slices (ROI: from the edge of the cerebral cortex until 250 nm from the edge of the cerebral cortex) after 30 min at the end of CM injection in coronal slices for sham group (blue, N = 5) and tDCS group (orange, N = 5). C. Representative images of CSF tracer distribution in coronal slices after 60 min at the end of CM injection. D. The mean intensity of CSF tracer in 8 slices (ROI: from the edge of the cerebral cortex until 250 nm from the edge of the cerebral cortex) after 60 min at the end of CM injection in coronal slices for sham group (blue, N = 5) and tDCS group (orange, N = 4). *p < 0.05, t-test

### tDCS influenced the efflux of ISF

The glymphatic system plays a pivotal role in brain health by facilitating the removal of metabolic wastes. Specifically, these wastes are cleared from the brain when the cerebral spinal fluid (CSF) exchanges with the interstitial fluid (ISF) and are subsequently excreted from the brain through various pathways, including the lymphatic system [27,28]. Given its properties, Direct Blue 53 (DB53), a small molecule, has been identified as a suitable tracer for efflux imaging [29]. In our study, to visualize and quantify the efflux of metabolic waste from the brain, we injected DB53 into the cisterna magna (CM) following tDCS stimulation (**Figs. 4A-B**). The distribution and concentration of DB53 were then observed microscopically in the cervical lymphatic vessels of the mice (**Figs. 4A-B**). Our observations between 30 to 60 minutes post-injection revealed that the concentration of DB53 in the tDCS group was significantly elevated compared to the sham group (**Figs. 4C-D**, Sham vs. tDCS, 139.06 ± 11.80 vs. 181.92 ± 12.07, p = 3.20E-2). This finding suggests that tDCS has a modulatory effect on the efflux of metabolic wastes from the mouse brain.

**Fig. 4.**
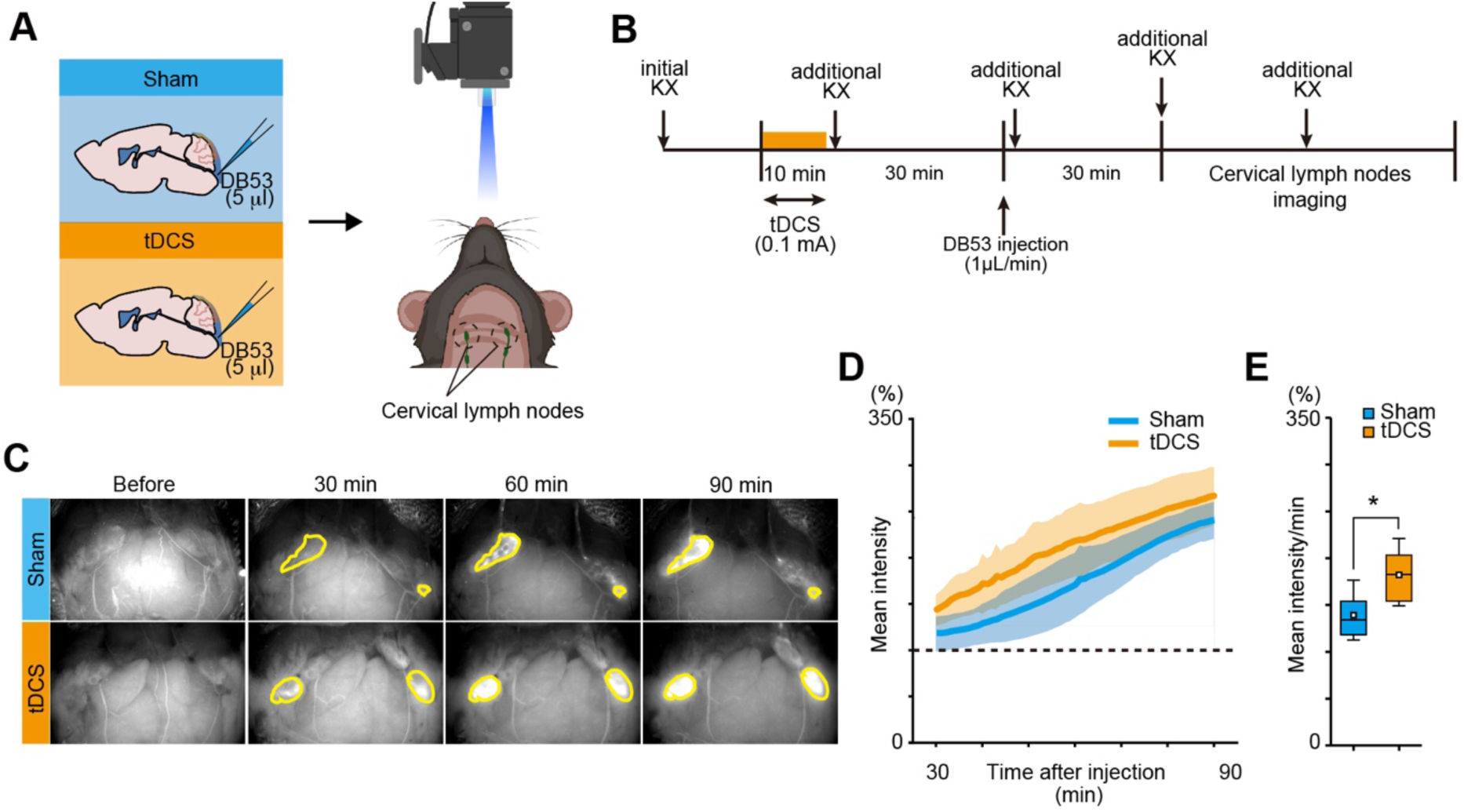
tDCS influenced the efflux of the glymphatic efflux. A-B. Schematic diagram for cervical lymph node imaging and the experiment schedule. C. Representative images of cervical lymph node imaging. D. Mean intensity (%) in cervical lymph nodes for sham group (blue, N = 5) and tDCS group (orange, N = 6). Shaded areas represent s.e.m. E. The rate of cervical lymph node filling of the first 30 min recording for sham group (blue, N = 5) and tDCS group (orange, N = 6). *p < 0.05, t-test

### No change in AQP4 expression in 30 min post-tDCS

Our findings suggest that tDCS affects the dynamics of influx and efflux within the mouse brain’s glymphatic system. Yet, the precise mechanism underlying this influence remains unclear. A pivotal element of the glymphatic system is the brain water channel, aquaporin-4 (AQP4). This protein is recognized for its crucial role in facilitating the exchange between CSF and ISF [22,30]. To ascertain whether tDCS impacts AQP4 expression, we utilized immunofluorescence staining techniques after tDCS administration and examined the expression patterns in the mouse brain (**Fig. 5A**). Specifically, we assessed the endfoot polarization of AQP4 expression. The perivascular polarity, defined as the ratio of the focally elevated perivascular signal area to the total AQP4 immunoreactivity (refer to Materials and Methods) [28], showed no significant difference between sham and tDCS-treated mice (**Figs. 5B-C**, Perivascular: Sham vs. tDCS, 139.43 ± 8.52 vs. 120.12 ± 7.29, p = 0.13). Similarly, the neuropil polarity, which represents the ratio of the neuropil signal area to the overall AQP4 immunoreactivity, also remained unchanged between the two groups (**Figs. 5B-C**, Neuropil: Sham vs. tDCS, 70.47 ± 6.71 vs. 70.95 ± 6.14, p = 0.96). These findings imply that the alterations in the glymphatic system’s dynamics induced by tDCS are not a result of changes in AQP4 expression, at least not within the 30-minute observation period.

**Fig. 5.**
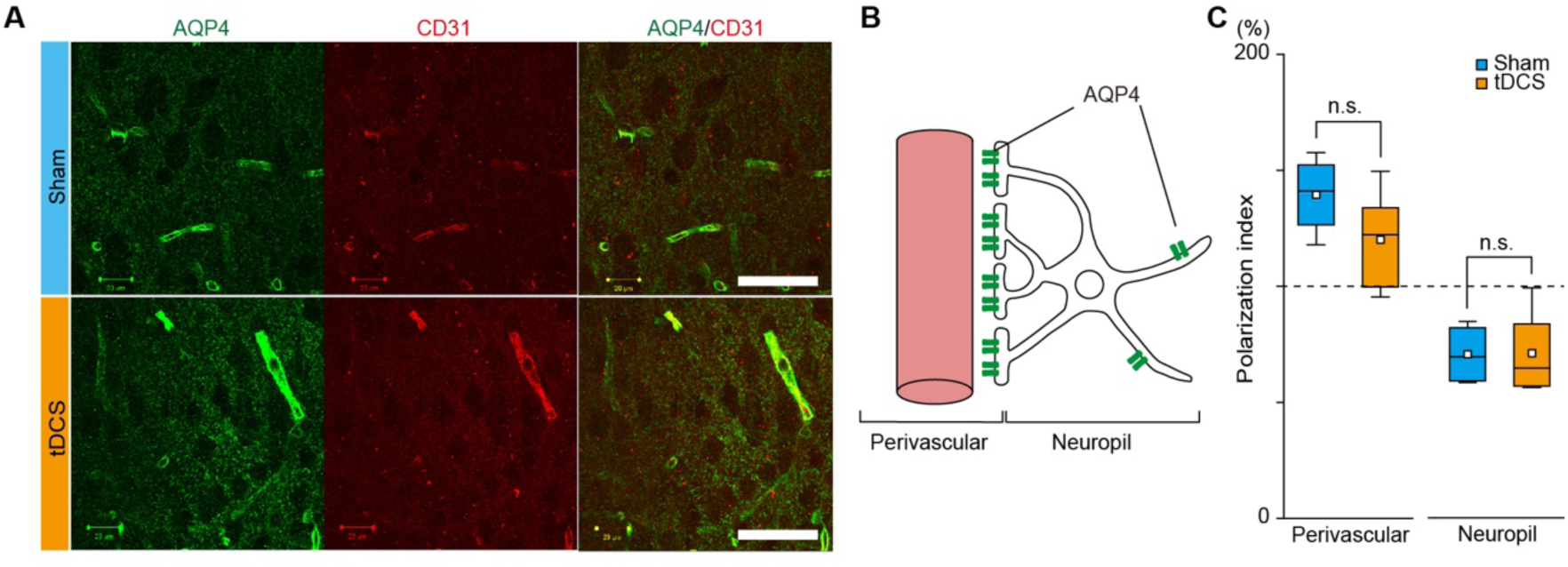
No change in AQP4/CD31 after tDCS. A. Representative images of AQP4/CD31 immunostaining. Scale bar: 50 *μ*m B. Schematic diagram of the position of perivalvular and neuropil for AQP4 polarization analysis. C. Polarization index (%) of AQP4/CD31 in perivascular and neuropil for sham group (blue, N = 4) and tDCS group (orange, N = 7).

### tDCS modulation of slow-wave sleep and its potential impact on brain clearance

Sleep is known to facilitate brain clearance by expanding the interstitial space during slow-wave sleep. This expansion results in enhanced CSF influx and efflux [20]. Given this understanding, we sought to determine if tDCS has any influence on slow-wave sleep, which could indirectly affect brain clearance. To this end, we conducted EEG recordings (**Fig. 6A**). Our findings revealed that, compared to the sham group, the delta wave amplitude increased 30 minutes post-tDCS. Additionally, the beta wave amplitudes significantly decreased 30 minutes post-tDCS (**Figs. 6B-C**, Delta wave (%): 30-60 min, Sham vs. tDCS, 100.71 ± 1.96 vs. 110.91 ± 5.44, p = 3.88E-2; 60-90 min, Sham vs. tDCS, 101.46 ± 3.85 vs. 115.81 ± 8.15, p = 7.78E-3; Beta wave (%): 30-60 min, Sham vs. tDCS, 102.80 ± 0.86 vs. 100.28 ± 1.86, p = 4.36E-2; 60-90 min, Sham vs. tDCS, 103.16 ± 1.05 vs. 100.07 ± 2.58, p = 8.19E-3). These EEG changes, particularly the augmentation of the delta wave, suggest that tDCS might modulate brain clearance, as delta wave activity is positively associated with this process.

**Fig. 6.**
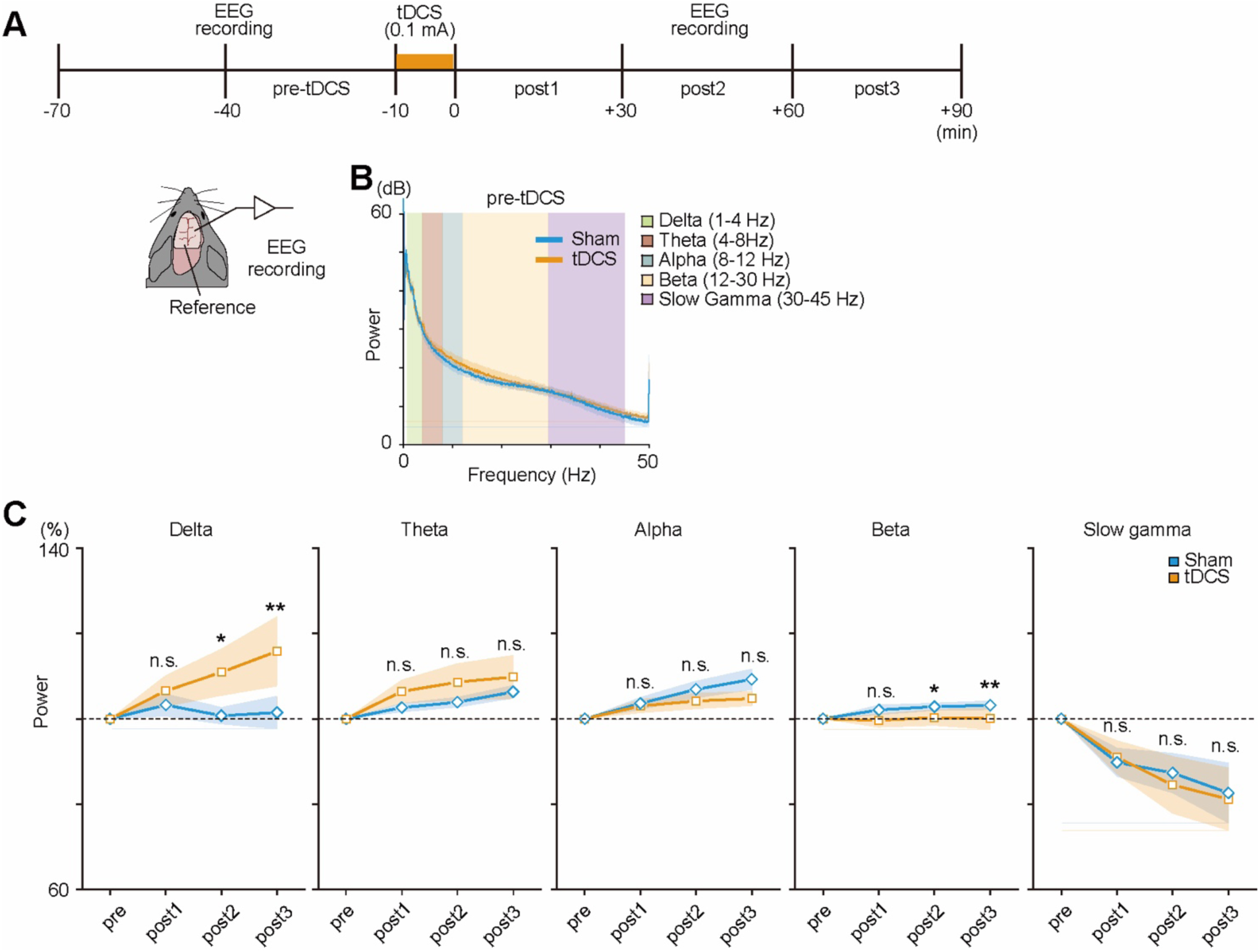
tDCS altered EEG power. A. Schematic diagram for EEG recordings and the experiment schedule. B. Normalized EEG power spectra for pre-tDCS period. C. Power (%) of delta wave (1-4 Hz), theta wave (4-8 Hz), alpha wave (8-12 Hz), beta wave (12-30 Hz), and slow gamma wave (30-45 Hz) in the sham group (blue, N = 5) and tDCS group (orange, N = 7) for different periods based on the baseline (-40∼-10 min). Shaded areas represent s.e.m. *p < 0.05, **p < 0.01, two-way repeated measures ANOVA

### tDCS effects on CSF-ISF exchange and brain waves absent in IP_3_R2-knockout mice

Previous studies have highlighted the relationship between delta wave activity and astrocytic functions. Specifically, Vaidyanathan et al. demonstrated that delta wave activity can be induced by astrocyte Ca^2+^ [31]. In a related study, Monai et al. identified that tDCS can trigger astrocyte IP_3_/Ca^2+^ signaling. Given these findings, we sought to determine if the observed increase in delta wave activity post-tDCS was indeed mediated by astrocyte IP_3_/Ca^2+^ signaling. To this end, we administered the CSF tracer, BDA, via the CM in the inositol trisphosphate receptor type 2-knockout (IP_3_R2 KO) mice (**Fig. 7A**). Again, our findings indicated no significant differences in CSF-ISF exchange between the sham and tDCS-treated groups (**Figs. 7B-C**, Cortex: Sham vs. tDCS, 59.51±5.55 vs. 64.20±5.47, p = 0.57; Hippocampus: Sham vs. tDCS, 66.40±8.54 vs. 74.84±4.21, p = 0.42; Dorsal striatum: sham vs. tDCS, 75.92±10.96 vs. 75.83±3.66, p = 0.99; Thalamus: sham vs. tDCS, 92.47±7.68 vs. 97.42±5.30, p = 0.62; Cerebellum: Sham vs. tDCS, 137.90±9.76 vs. 121.40±5.36, p = 0.20; Frontal: Sham vs. tDCS, 120.65±16.18 vs. 95.81±17.03, p = 0.32; Motor: Sham vs. tDCS, 51.40±5.21 vs. 54.55±5.99, p = 0.70; Somatosensory: Sham vs. tDCS, 55.25±7.99 vs. 60.77±4.61, p = 0.58; Visual: Sham vs. tDCS, 60.29±13.46 vs. 68.78±7.29, p = 0.60). Subsequently, we conducted EEG recordings on IP_3_R2 KO mice (**Fig. 7E**). Our results showed no significant alterations in delta wave, beta wave (**Fig. 7F**, Delta wave (%): 0-30 min, Sham vs. tDCS, 85.09±5.42 vs. 95.38±5.30, p = 0.29; 30-60 min, Sham vs. tDCS, 83.82±5.98 vs. 92.95±4.57, p = 0.34; 60-90 min, Sham vs. tDCS, 76.63±9.15 vs. 89.58±6.30, p = 0.20; Beta wave (%): 0-30 min, Sham vs. tDCS, 114.72±5.15 vs. 105.50±4.62, p = 0.28; 30-60 min, Sham vs. tDCS, 115.64±5.63 vs. 106.33±4.94, p = 0.27; 60-90 min, Sham vs. tDCS, 117.86±8.34 vs. 108.12±6.77, p = 0.32) and other brain wave activities (Theta wave (%): 0-30 min, Sham vs. tDCS, 67.65±12.31 vs. 67.97±7.90, p = 0.95; 30-60 min, Sham vs. tDCS, 68.21±13.77 vs. 65.33±6.62, p = 0.91; 60-90 min, Sham vs. tDCS, 56.16±13.27 vs. 67.87±9.17, p = 0.38; Alpha wave (%): 0-30 min, Sham vs. tDCS, 153.39±32.55 vs. 158.12±51.26, p = 0.79; 30-60 min, Sham vs. tDCS, 151.95±33.85 vs. 158.33±48.56, p = 0.80; 60-90 min, Sham vs. tDCS, 178.94±35.43 vs. 109.49±18.47, p = 0.16; Slow gamma wave (%): 0-30 min, Sham vs. tDCS, 99.97±1.86 vs. 106.76±3.70, p = 9.85E-2; 30-60 min, Sham vs. tDCS, 100.69±1.39 vs. 107.77±3.51, p = 8.93E-2; 60-90 min, Sham vs. tDCS, 103.38±2.03 vs. 111.07±4.63, p = 9.29E-2) in these mice. Based on these observations, we conclude that the IP_3_/Ca^2+^ pathway plays a crucial role in the tDCS-induced increase in delta wave activity and decrease beta wave activity.

**Fig. 7.**
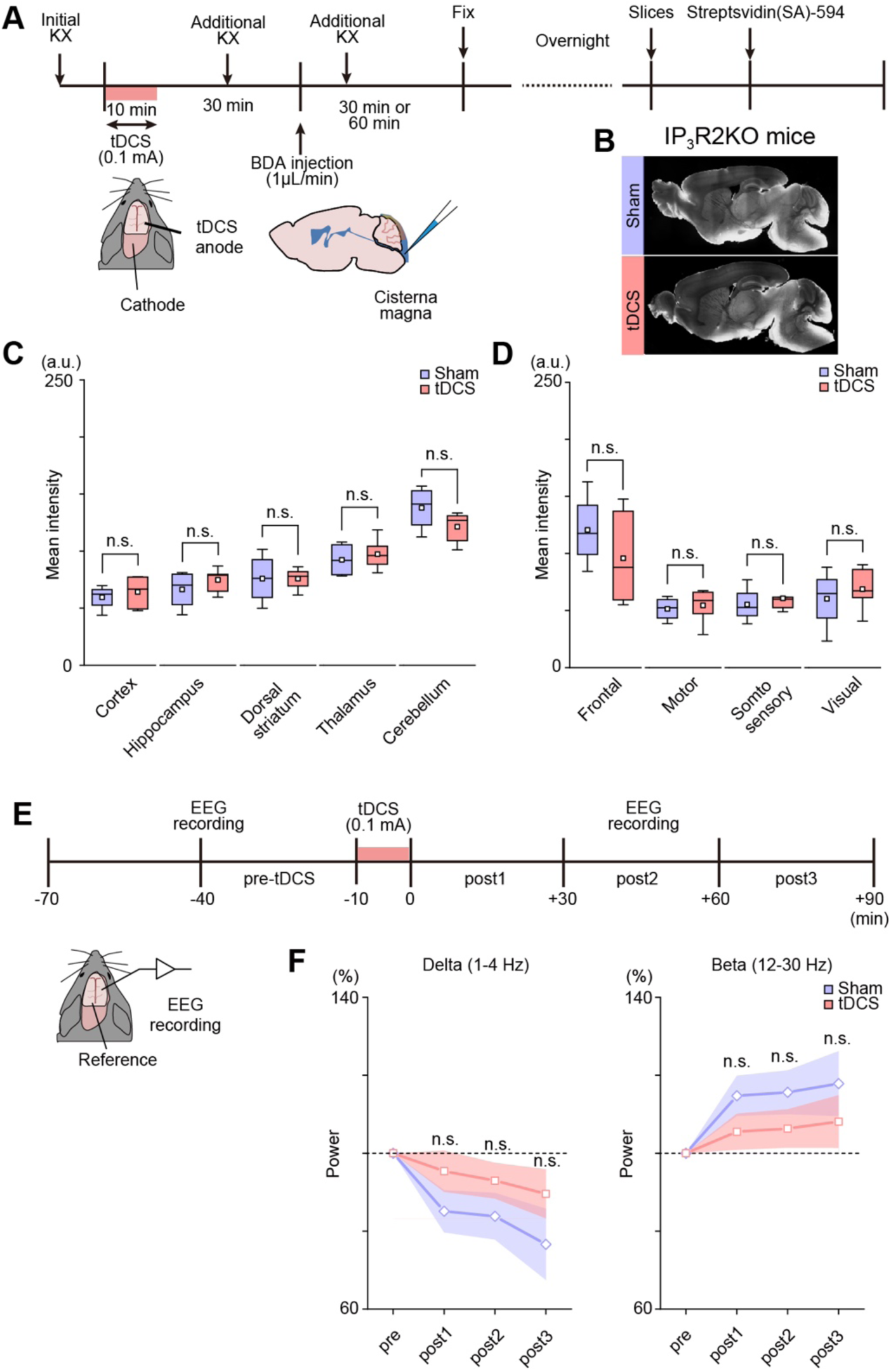
IP_3_/Ca^2+^ signaling plays an essential role in tDCS-induced CSF influx. A. Schematic diagram for tDCS and the experiment schedule. B. Representative images of CSF tracer distribution in IP_3_R2-KO mice’s sagittal slices. C-D. The mean intensity of the CSF tracer after 30 min at the end of CM injection in sham group (purple, N = 4) and tDCS group (pink, N = 6) for IP_3_R2-KO mice’s sagittal slices. E. Schematic diagram for EEG recordings and the experiment schedule. F. Power (%) of delta (1-4 Hz) and beta wave (12-30 Hz) in sham group (purple, N = 5) and tDCS group (pink, N = 6) for IP_3_R2-KO mice. Shaded areas represent s.e.m.

## Discussion

In this study, we observed an increase in CSF tracer in the mouse brain after tDCS by using the CSF tracer method, suggesting that tDCS affects CSF influx and thus alters CSF-ISF exchange.

The mechanism of tDCS as a non-invasive brain stimulus is very complex, involving changes in various neurotransmitters, including dopamine, acetylcholine, and serotonin. Mishima et al. suggested that tDCS may enlarge microglial somata by inducing noradrenergic signaling in awake mice [32]. Monai et al. found that for the cellular mechanisms of anodal tDCS, the activation of adrenergic receptors plays a significant role in the mouse brain after tDCS [33]. These results relate to noradrenaline, which facilitates the transition from the sleep state to the awake state, which is essential for the clearance of metabolic waste from the brain [34]. However, all experiments in this study were conducted under anesthesia, and the anesthetic was replenished at 30-minute intervals to ensure that the animals were adequately anesthetized, leading us to believe that the increase in noradrenaline did not alter the anesthetic state of the animals.

According to the glymphatic system, CSF, after flowing into the brain, exchanges with ISF in the extracellular space, carrying metabolic waste out from the ISF of the brain. It has been shown that the extracellular space becomes larger after tDCS, which facilitates the diffusion of solutes in the brain [35]. The extracellular space enlarges, and as a result, CSF flows into this expanded extracellular space, thus accelerating the CSF-ISF exchange, which further supports our results.

CSF carries brain metabolic wastes out of the brain after the exchange with ISF. There are three pathways for the efflux of metabolic waste from the brain: (1) arachnoid granulations; (2) meningeal lymphatics; (3) and nasal lymphatics parenchyma. The first of these pathways is controversial, and the second and third both pass through the cervical lymph nodes [27]. Therefore, in this study, we injected the CSF tracer into the cisterna magna (CM) and then conducted cervical lymph nodes imaging to monitor the efflux of metabolic waste from the brain. We observed an increase in the rate of CSF tracer efflux from the cervical lymph nodes after tDCS via cervical lymph node imaging, suggesting that tDCS alters the CSF-ISF exchange and affects the efflux of the glymphatic system.

The significant decrease in glymphatic clearance in AQP4-knockout mice suggests an essential role for AQP4 in the glymphatic system [23]. However, we did not find changes in A after tDCS in this study, although Luo et al. found that AQP4 expression was enhanced in AD model mice after two months of daily tDCS for 30 min [17]. Since tDCS was only performed once, and the brain was examined 30 min after stimulation, we plan to extend the treatment period of tDCS in future experiments.

Xie et al. found that the CSF-ISF exchange in the sleep and anesthesia state is much more enhanced than in the awake state since, due to a 60% increase in interstitial space, which is related to the increased delta wave that enlarges the interstitial space [21]. In this study, we found that the delta wave increased 30 minutes after tDCS. This result suggested that an increased delta wave is one of the reasons that tDCS enhances the CSF-ISF exchange. Additionally, we also found that the beta wave decreased 60 minutes after tDCS stimulation. Hablitz et al. found that the enhancement of brain clearance was positively correlated with the delta wave and negatively correlated with the beta wave [36], which is consistent with the results of this study.

Astrocytes have long been recognized as the supportive cells of the central nervous system; with the continuous advancement of research, it has become increasingly evident that astrocytes play an important role in various aspects of neurobiology[37]. Regarding the mechanism of tDCS, Monai et al. have found that tDCS could induce astrocytic IP_3_/Ca^2+^ signaling [33]. This astrocytic intracellular Ca^2+^ signal can cause a series of changes, including altering neuronal information processing, extracellular ion concentration, and more [38]. Additionally, numerous studies have demonstrated that the anodal electrode of tDCS induces a rise in cerebral blood flow[39], which is influenced by the parameters and duration of tDCS[40–42]. However, recent research indicates that

tDCS-induced blood flow augmentation is not associated with IP_3_/Ca^2+^ signaling[43]. In our study, there were no significant differences observed in CSF-ISF exchange or brain wave patterns, particularly delta and beta waves, in IP_3_R2 KO mice. Therefore, in our study, the enhancement of tDCS-induced brain clearance in wild-type mice can be attributed to an increase in delta waves mediated by IP_3_/Ca^2+^ signaling, and we will conduct further research on the molecular mechanism in the future.

## Conclusions

In conclusion, we found that tDCS enhanced the influx and efflux of the glymphatic system, although there was no significant change in A expression. However, we found through the astrocyte IP_3_/Ca^2+^ signaling, tDCS alters the delta wave, which is positively correlated with brain clearance. These results suggested that tDCS altered the clearance of brain metabolic waste, providing theoretical support for the clinical application of tDCS.

## Acknowledgement

This work was supported by Ochanomizu University, KAKENHI grants (18K14859, 20K15895), JST FOREST Program, Grant Number JPMJFR204G, Research Foundation for Opto-Science and Technology, Kao Research Council for the Study of Healthcare Science, The Japan Association for Chemical Innovation, and TERUMO LIFE SCIENCE FOUNDATION.

